# A Single Population Approach to Modeling Growth and Decay in Batch Bioreactors

**DOI:** 10.1101/2025.04.27.650866

**Authors:** Carlos Martínez, Marielle Péré

**Affiliations:** Escuela de Ingeniería Bioquímica, Pontificia Universidad Católica de Valparaíso, Av. Brasil 2085, Valparaíso, Chile; Université Côte d’Azur, Inria, INRAE, CNRS, Sorbonne Université MACBES Team, Sophia-Antipolis, France; Université Côte d’Azur, CNRS UMR 7275, Inserm 1323, Institut de Pharmacologie Moléculaire et Cellulaire, Sophia-Antipolis, France

**Keywords:** maintenance rate, bioprocess, integro-differential equation, mortality rate, parameter estimation, kinetic modeling

## Abstract

Fitting dynamic models to population data such as the logistic growth equation is a common practice for describing microbial growth, both in natural ecosystems and under research conditions. However, these models are limited to scenarios where the population stabilizes at equilibrium, making them unsuitable for systems like batch bioreactors, where populations decline after substrate depletion. In this work, we propose two new population models capable of accurately describing such dynamics while maintaining an interpretable structure in which each parameter has a biological meaning. These models incorporate the growth rate as a function of cumulative biomass, rather than solely the biomass concentration, thereby accounting for memory effects. We establish fundamental properties of these models and demonstrate their applicability and accuracy to describe data from batch bioreactors.

## 1 Introduction

Mathematical models of population dynamics play a fundamental role in describing microbial growth and predicting biological behaviors in bioprocesses. Among these, the logistic growth model is one of the most widely used models due to its ability to capture sigmoidal growth patterns in a simple and intuitive manner [1]. This model has been extensively applied, for example, in studies on microalgae [2, 3], bacteria [4, 5, 6], and yeast [7]. However, a limitation of the logistic growth model is its inability to account for the decline phase observed in batch cultures. A batch culture is a closed system where microorganisms grow in a fixed volume of nutrient medium without additional inputs or removal of culture components. Typically, microbial growth in batch systems follows three main phases [8]: an initial exponential growth phase, a stationary phase, and a subsequent decay phase. While logistic growth models effectively describe the exponential and stationary phases, they fall short in capturing the subsequent decay phase that follows resource depletion. This limitation is significant, as the decay phase provides insights into energy costs associated with cell maintenance. For instance, in mammalian cell cultures, cell density often declines sharply after the stationary phase [9], while in microalgal cultures, a similar decline occurs due to maintenance energy demands and respiration [10]. Because the standard logistic growth equation assumes a fixed carrying capacity without a mechanism for population decline, it is insufficient for capturing the full dynamics of such biological systems.

Some variations of the single-population logistic growth model have been proposed to incorporate a decline phase [11, 12, 13]. For example, the Volterra integro-differential equation (IDE) [11]:

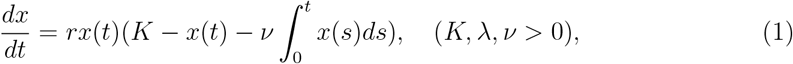

where *r* represents the intrinsic growth rate and *K* the carrying capacity. The additional integral term accounts for the cumulative effects of metabolism or toxin accumulation over time, which contribute to population decline. Here, *ν* acts as a sensitivity factor regulating how strongly these accumulated effects influence mortality. When *ν* = 0, this equation reduces to the classic logistic growth model. Other generalizations rely on purely descriptive functions without providing an explicit mechanistic explanation for the decline phase. For instance, [14] proposed the following empirical model:

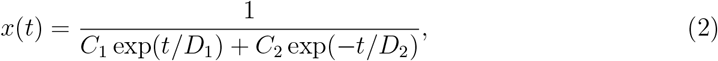

with *C*_1_, *C*_2_, *D*_1_, *D*_2_ non-negative parameters. Additionally, some studies have considered time-dependent parameter models, where growth and death rates evolve dynamically to reflect physiological changes in the population [15]. While some of these models are able to accurately describe population decline [16, 17], their parameters are not easily linked to the kinetic parameters of batch reactors. This limitation reduces their applicability beyond specific experimental conditions, making it challenging to generalize their use in broader bioprocess contexts.

The objective of this study is to develop single-population models capable of describing both the growth and decay phases observed in batch bioreactors while maintaining biologically meaningful parameters. To achieve this, we build upon classical models of microbial growth in batch cultures, identifying key limitations in existing approaches that fail to capture population decline [18, 19]. By introducing appropriate assumptions on growth kinetics, we derive two new single-population models formulated as integro-differential equations. These models not only improve the representation of microbial dynamics in batch systems but also extend their applicability to any biological process where population decline is a crucial factor. Furthermore, to facilitate numerical implementation and parameter estimation, we demonstrate that our models can be reformulated as a system of ordinary differential equations (ODEs), allowing for straightforward integration using standard computational techniques.

The paper is organized as follows. In Section 2, we revisit the equations for modeling batch cultures and derive single-population growth models. Section 3 introduces a generalized single-population model, together with a discussion of its fundamental properties. In Section 4, we validate the proposed models using two distinct datasets: one from a mammalian cell culture and another from a bacterial culture. Finally, Section 5 presents a discussion of our findings and their implications.

## 2 Derivation of the population models

The dynamics of microbial growth and substrate consumption in batch cultures can be mathematically described using a system of ODEs as follows [18, 19]:

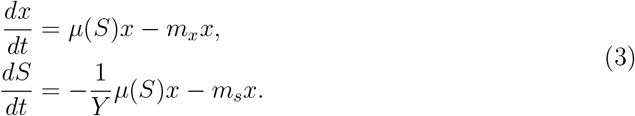

Here, *x* represents the population biomass concentration (e.g., in g m^−3^), *S* denotes the substrate concentration (e.g., in g m^−3^), *µ*(*S*) is the specific growth rate (e.g., in h^−1^), and *Y* is the apparent yield coefficient (e.g., in g g^−1^). The parameters *m*_*x*_ and *m*_*s*_ are associated with the maintenance of the population. The term *m*_*x*_ is the specific maintenance rate (e.g., in h^−1^), and may be regarded as an endogenous metabolism rate resulting in the consumption of maintenance energy and a decrease in the biomass, while *m*_*S*_ is associated with the consumption of substrate for non-growth functions [20]. Model (3) is complemented with *x*_0_ and *S*_0_ as the initial biomass and substrate concentrations, respectively. That is

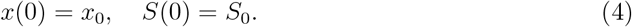

The inclusion of maintenance terms is crucial to accurately capture the decline phase. As resources are depleted, the specific growth rate *µ* eventually drops to zero, and population dynamics transition into a decay phase driven by maintenance energy demands. Without maintenance terms, any solution to (3) would lead to a stationary biomass concentration, failing to reflect the eventual decline observed in real batch cultures.

Various models have been proposed in the literature to describe the specific growth rate *µ* [21]. The most widely used approach is Monod kinetics, which provides an empirical description of microbial growth [22]. However, alternative models are often adopted to enhance predictive accuracy [23]. In this study, our focus is on deriving a single-population model from (3), which requires growth rate formulations that facilitate its mathematical analysis. One such alternative is the Blackman model [21] (see Figure 1), which is valid for low substrate concentrations and reads

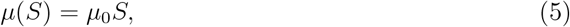

**Figure 1:**
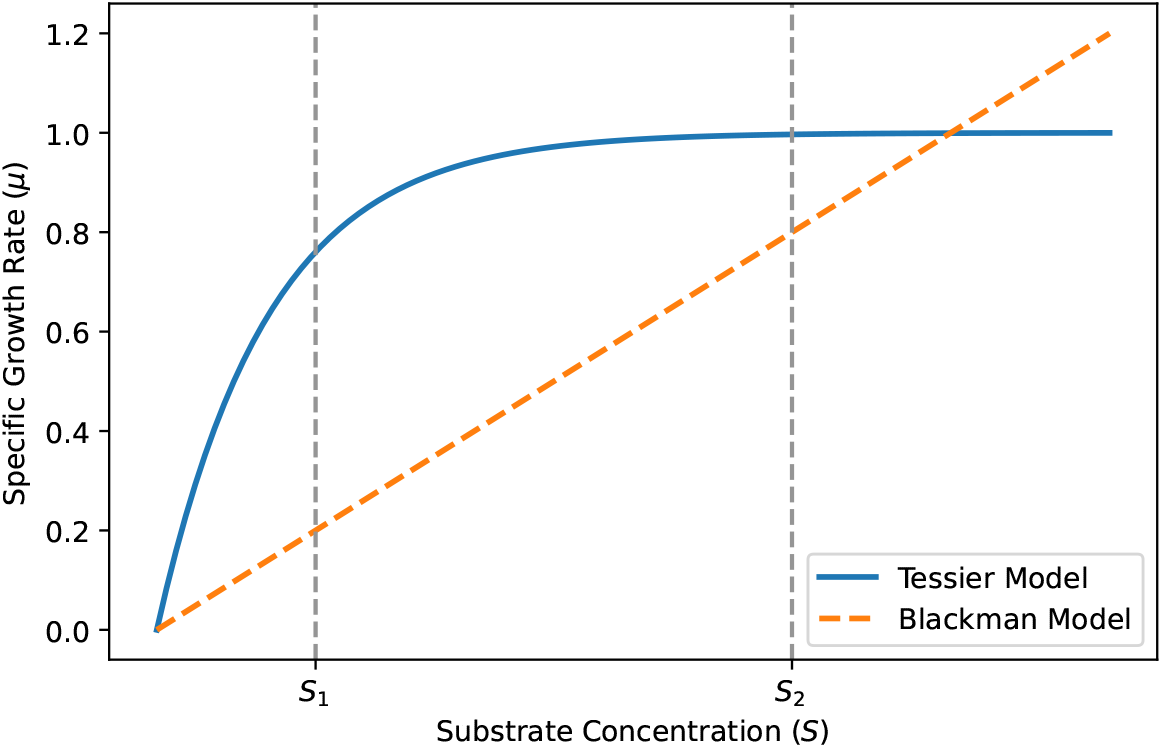
Comparison of the specific growth rate functions given by (5) and (6). Parameters are chosen as *µ*_*max*_ = 1, *K*_*s*_ = 0.7, and *µ*_0_ = 0.2.

with *µ*_0_ a proportionality constant. Another alternative is the Teissier model [21] (see Figure 1), given by

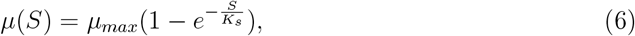

with μ_*max*_ the maximum specific growth rate, and *K*_*s*_ a saturation constant.

In the following, we derive single-population models from (3) by substituting *µ* with the Blackman model (5) and the Teissier model (6). The Blackman model is selected for its simplicity, and also because in absence of maintenance terms, (3) reduces to the classical logistic growth equation [24]. The Teissier model is selected because it provides a bounded growth function, unlike the Blackman model, which assumes linear dependence. This makes it more appropriate for conditions where resource availability is initially abundant. Additionally, the Teissier function captures the diminishing returns effect observed in many biological systems, where increasing substrate concentrations lead to progressively smaller increases in growth rate (see Figure 1).

### 2.1 Single population model based on Blackman model

When *µ* is described by (5), the batch model (3) becomes

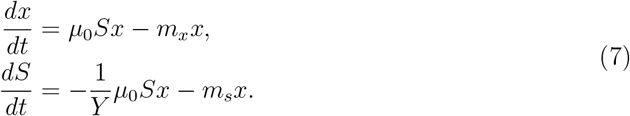

While the batch model (7) provides a direct description of biomass and substrate dynamics, it can be reformulated in a more compact form that explicitly accounts for cumulative biomass effects. This leads to the following IDE equation, which we establish in the next proposition:

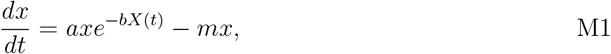

with *a, b*, and *m* non-negative parameters, and *X*(*t*) the cumulative biomass (or cumulative population) defined as:

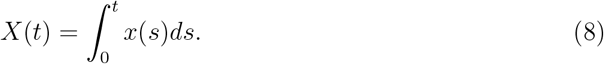

#### Proposition 2.1.

*Let* (*x, S*) *be a solution of (4)-(7), then x solves model M1. Moreover, we have that*

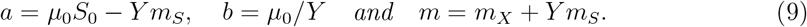

*Proof*. From the second equation in (7), we have that

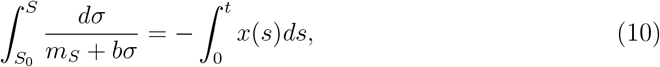

with 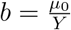. Integrating (10), we obtain

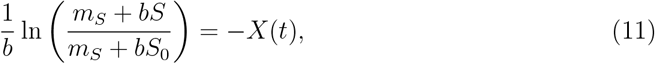

This implies that

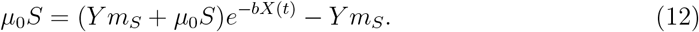

Now, replacing *μ*(*S*) in (7) by (12), we complete the proof. □

The formulation of model M1 captures the dependence of biomass growth not only on its current concentration but also on its cumulative history, reflecting the depletion of resources over time. The growth term introduces a memory effect, where the integral of biomass accounts for past abundance: higher cumulative biomass implies greater resource consumption, reducing future growth rates. Parameter *a* represents growth rate while *b* is related to depletion of nutrients. Specifically, for large values of *b*, the growth rate rapidly approaches zero as the population grows, leading to faster decline.

Model M1 depends on three parameters, *a, b*, and *m*, whereas the original formulation (7) depends on four. This shows the impossibility of estimating all the parameters of (7) based only on population data (data for *x*).

### 2.2 Single population model based on Teissier model

When *µ* is given by (6), then (3) becomes

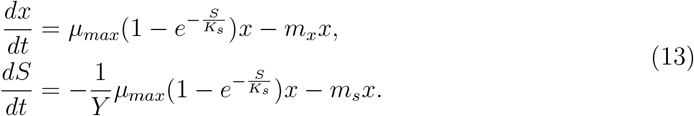

In the following proposition, we prove that the population *x* satisfies the following IDE, which we will refer to as model M2:

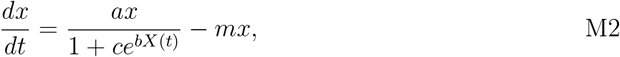

with *a, b, c*, and *m* non-negative parameters, and *X*(*t*) defined by (8).

#### Proposition 2.2.

*Let* (*x, S*) *be a solution of (4)-(13), then x solves model M2. Moreover*,

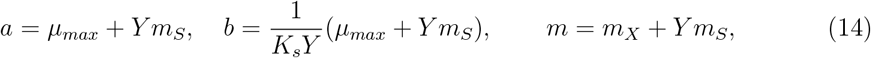

*and*

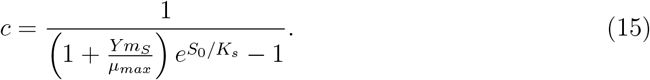

*Proof*. From the second equation in (13) we obtain that

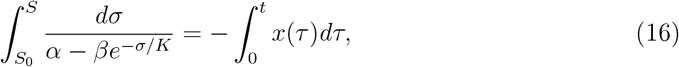

with 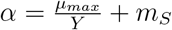 and 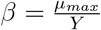. Integrating (16), we obtain that

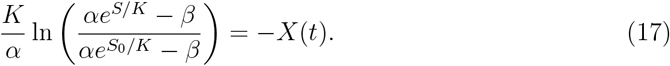

This implies that

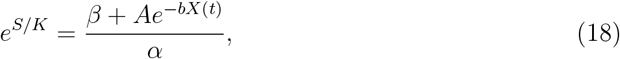

with 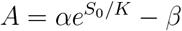 and 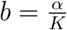. Now, reordering (18), we obtain that

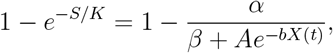

thus, we conclude that

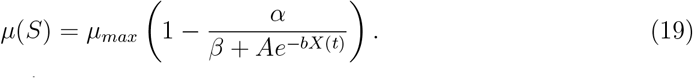

Since *β* − *α* = −*m*_*S*_, we can write

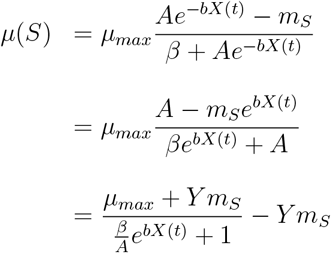

Replacing (19) in the first equation of (13) we get

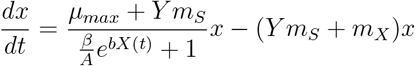

Finally, defining *a* = *µ*_*max*_ + *Y m*_*S*_, *c* = *β/A* and *m* = *m*_*X*_ + *Y m*_*S*_, we complete the proof. □

Model M2 depends on four parameters (*a, b, c*, and *m*), while the batch model (13) depends on five parameters. This demonstrates that it is impossible to estimate all parameters of (13) using only population data (i.e., measurements of *x*). In this case, parameter *a* represents the net growth rate of the population, while *b* determines the rate of resource depletion. The parameter *c* accounts for the abundance of resources. Indeed, according to (15), as the initial concentration of substrate *S*_0_ increases, the parameter *c* becomes closer to zero.

Note that if *m*_*S*_ = 0, which is a common assumption in batch culture models, then the meaning of each parameter becomes more straightforward. Indeed, from (14) and (15), we obtain that

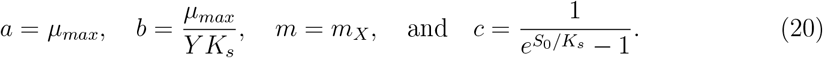

Expression (20) shows a direct interpretation of the model parameters. Specifically, *a* corresponds to the maximum specific growth rate, and *m* to the specific maintenance rate. The half-saturation constant can be determined from *c* and the initial substrate concentration:

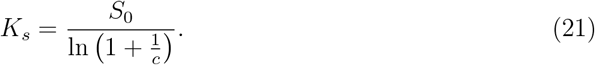

Finally, *Y* can be determined from *b*.

## 3 Properties of the models

### 3.1 Generalized single-population model representation

Both models proposed in the previous section, M1 and M2, can be written in the following form:

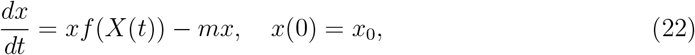

where *x* is the biomass concentration, *m* is a term accounting for the decline of the population (e.g., due to maintenance or mortality), *X*(*t*) is the cumulative biomass given by (8), *x*_0_ *>* 0 is the initial biomass concentration, and *f* is a function representing the growth of the population, which takes one of the following forms:

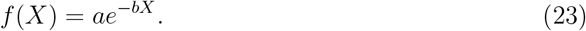

or

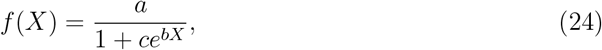

The choice between (23) and (24) depends on the available data and the specific objectives of the study. Since (24) was originally derived from the Teissier model, this function is theoretically better suited for scenarios where resources are initially abundant and the maximum specific growth rate is reached. On the other hand, since (23) was originally based on the Blackman model, this function is more appropriate when resource availability initially limits growth.

### 3.2 Comparing population models

We observe that in both models, M1 and M2, when biomass is cumulated the dominant term exhibits an exponential decrease dependent on *m*. Thus, if both models describe the same population, they should share a similar value of *m*. Therefore, to facilitate comparison of both models, we introduce the dimensionless time variable *τ* = *mt* and define the normalized biomass *y*(*τ*) = *x*(*t*)*/x*_0_. This leads to the following dimensionless formulations of M1 and M2, denoted as M1d and M2d, respectively:

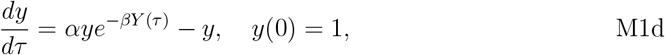

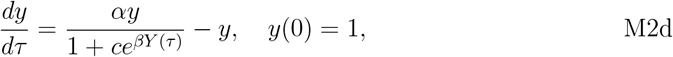

where 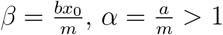, and 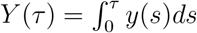.

Figure 2 presents simulations of M1d and M2d. For low values of *c*, model M2d captures a rapid initial population increase, a behavior that M1d fails to reproduce (see Figure 2A). Based on the batch models from the previous section, this can be explained as follows. Low values of *c* imply high initial substrate concentration in the bioreactor (see (15)). Now, since Model M1d is based on the Blackman model (5), any reduction in substrate concentration immediately leads to a decrease in the growth rate. In contrast, since M2d is based on the Teissier model (6), it offers more flexibility and can describe situations where the growth rate remains maximum over a period, even as substrate levels decrease. This is illustrated in Figure 1. If the initial substrate concentration is *S*_2_, then an initial consumption of substrate has a different impact on each specific growth rate model.

**Figure 2:**
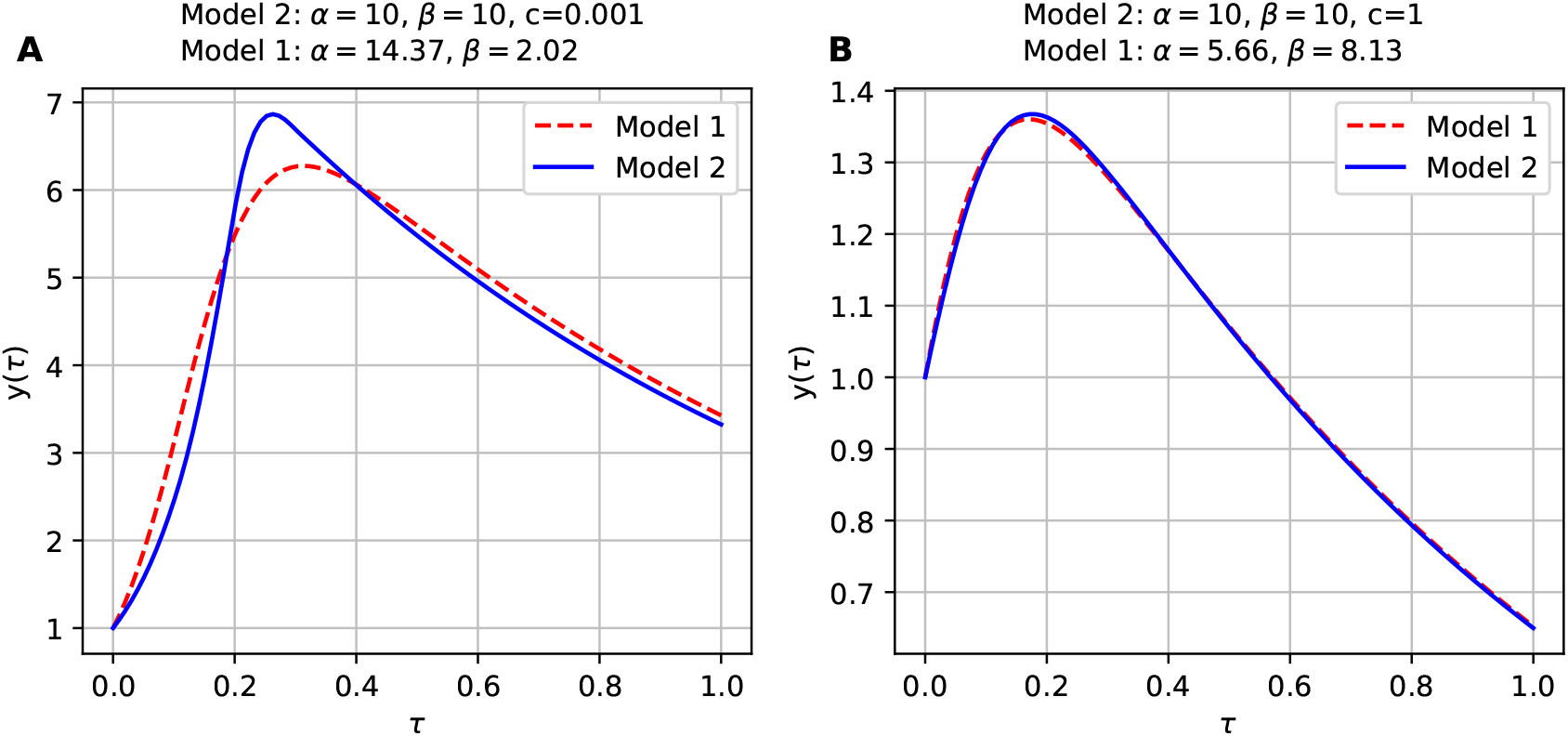
Simulations of the dimensionless models M1d and M1d for different sets of parameters.

Figure 2B shows that both models can yield similar results. In such a case, a larger value of *c* is considered. Following expression (14), larger values of *c* are associated with lower initial substrate concentrations. As shown in Figure 1, for example at concentration *S*_1_, both specific growth rate models have a similar behavior, in the sense that both functions have a notorious decrease as the substrate concentration decreases.

### 3.3 Numerical implementation

A key property of any mathematical model is its numerical solvability. There is a vast body of literature on numerical methods for solving IDEs, and several studies have specifically addressed numerical approaches for single population models such as the Volterra equation (1) [25, 26]. However, the mathematical technicalities of these methods may discourage the use of IDEs in applied fields such as bioprocess modeling. Therefore, we propose to define the auxiliary variable *z*(*t*) = *X*(*t*), to rewrite (22) as the following system of ordinary differential equations (ODEs)

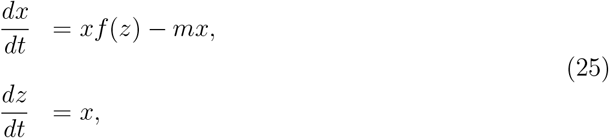

with the initial conditions *x*(0) = *x*_0_ and *z*(0) = 0. Solving (25) is straightforward using existing numerical methods for ODEs.

### 3.4 Long-term behavior

We show that for a broad class of functions *f*, any solution to (22) will ultimately decline to zero. Thus, this kind of population model is not suitable for populations that stabilize at a positive equilibrium.

#### Proposition 3.1.

*Consider (22) and let us assume that f is strictly decreasing and that* lim_*X*→∞_ *f* (*X*) = 0. *Then any solution x*(*t*) *to (22) satisfies x*(*t*) → 0 *as t* → ∞.

*Proof*. Let us assume that *f* (0) *< m*. Since *f* is strictly decreasing, we have that

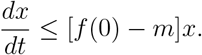

Using Grönwall’s inequality, we obtain

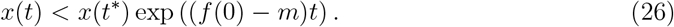

Since *f* (0) *< m*, from (26), we conclude that *x*(*t*) → 0 as *t* → ∞. Now, let us assume that *f* (0) ≥ *m*. Since *f* is strictly increasing and lim_*x*→∞_ *f* (*x*) = 0, there exists *X*^*^ ≥ 0 such that *f* (*X*^*^) = *m*. Now let us assume that

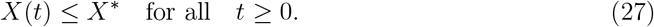

In that case, *f* (*X*(*t*)) ≥ *m* for all *t* ≥ 0, which implies 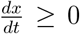. Thus, *x*(*t*) ≥ *x*_0_ *>* 0 for all *t* ≥ 0. Using the definition of *X*(*t*) (8), we obtain that *X*(*t*) ≥ *x*_0_*t*, contradicting the (27). Therefore, there exists *t*^*^ *>* 0 such that *X*(*t*^*^) *> X*^*^. Since 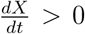, it follows that *X*(*t*) *> X*^*^ for all *t > t*^*^, and consequently

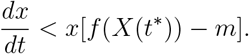

Using Grönwall’s inequality, we obtain

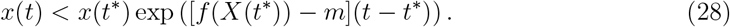

Since *f* (*X*(*t*^*^)) −*m <* 0, from (28), we conclude that *x*(*t*) → 0 as *t*→ ∞. This completes the proof. □

## 4 Fitting observed data

To assess the ability of models M1 and M2 to accurately represent data, we consider two different datasets from the literature. The first one corresponds to data from a mammalian cell culture provided by [16], and the second one corresponds to a culture of the bacterium *Lactobacillus helveticus* provided by [27]. We simulate the models (M1) and (M2) using their formulation as a system of ODEs (25). The integration is performed with a Runge-Kutta method, specifically the Dormand-Prince method (RK45), implemented in Python via the `solve_ivp` function from the `SciPy` library. The corresponding Python code is available at https://github.com/mariellePr/Extended-Logistic-Growth-Model.

### 4.1 Growth of mammalian cells

Mammalian cell cultures can be used for the production of therapeutic proteins, vaccines, and advancing biomedical research by mimicking complex cellular environments [28]. In batch cultures, mammalian cells undergo a sharp decline in density after the stationary phase, making this system a good candidate for testing models M1 and M2. We consider data from two mammalian cell cultures provided by [16]. As indicated in the caption of Figure 3, we will refer to these datasets as A and B.

**Figure 3:**
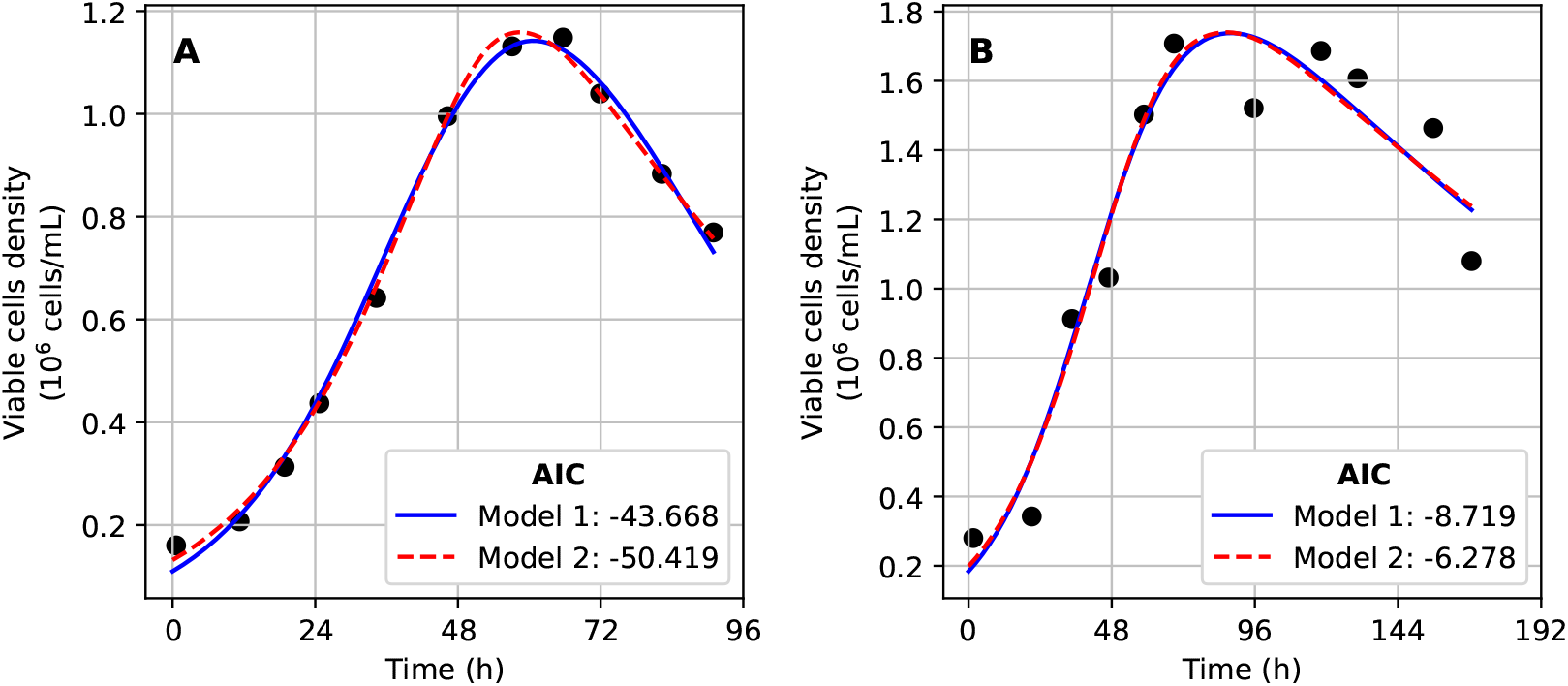
Model calibration using two sets of mammalian cell culture data presented in [16]. The blue line represents the solution of model M1, while the dashed red line corresponds to M2.The black dots correspond to data. Table 1 summarizes the parameter values. (A) Models fitted to Dataset A. (B) Models fitted to Dataset B.

To estimate model parameters, we solve a least-squares optimization problem for each dataset *j* ∈ {*A, B*} and each model *M* ∈ {*M* 1, *M* 2}. Each model *M* is governed by a parameter set 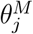, which must be fitted separately for each dataset. Given observations at non-uniform time points 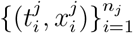, the optimal parameters 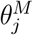 minimize the squared error:

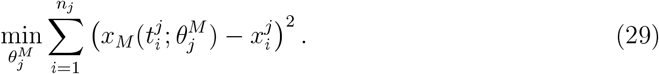

**Table 1:** Estimated parameters for model M1 and M2 when they are fitted to the datasets from Figure 3.

| Dataset | Model | $a$ ($\text{h}^{-1}$) | $b$ ($\text{g}^{-1} \text{L h}^{-1}$) | $c$ | $m$ ($\text{h}^{-1}$) | AIC |
| --- | --- | --- | --- | --- | --- | --- |
| Dataset A | M1 | 0.122 | $0.019 \times 10^{-6}$ | - | 0.059 | -43.668 |
| | M2 | 0.074 | $0.104 \times 10^{-6}$ | 0.077 | 0.019 | -50.419 |
| Dataset B | M1 | 1.381 | $0.582 \times 10^{-6}$ | - | 0.146 | -8.719 |
| | M2 | 2.692 | $0.755 \times 10^{-6}$ | 1.159 | 0.130 | -6.278 |

Here, 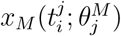 represents predictions by model *M* at time 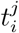 given parameters 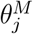, ensuring the estimation accounts for dataset-specific sampling points. To solve (29), we employ the Levenberg-Marquardt algorithm using the `curve_fit` function from the `SciPy.optimize` [29] module in Python.

Additionally, we assess the performance of both models using the Akaike Information Criterion (AIC) [30], which balances goodness of fit and model complexity. Lower AIC values indicate a better trade-off between model accuracy and complexity.

Figure 3 presents the results, showing that both models, M1 and M2, accurately describe the data. However, as seen in Figure 3A, model M2 achieves a lower AIC than M1, despite its higher complexity. In contrast, Figure 3B shows that both models yield nearly identical curves, but in this case, M1 has a slightly lower AIC. Although the AIC values are close, model selection can also be guided by the interpretability of parameters. As discussed in Section 2, the parameter *a* represents the specific growth rate, but its interpretation depends on the model. In Model M1, *a* represents the initial growth rate given the starting substrate conditions, whereas in Model M2, *a* corresponds to the maximum achievable growth rate, independent of initial conditions. This distinction highlights the flexibility of Model M2 in capturing prolonged exponential growth under nutrient-rich environments. This model should be used in case that the maximum specific growth rate is reached under initial conditions. Parameter *m* is associated with the specific mortality rate.

### 4.2 Growth of *Lactobacillus helveticus*

We analyze the data presented in [27] (see Figure 4), which represents the biomass dynamics of the bacterium *Lactobacillus helveticus* under different cultivation conditions. Each curve in Figure 4 represents a distinct condition (*j* = 1, 2, 3, 4), which is characterized by a different initial concentration of yeast extract, a nutrient that influences bacterial growth and metabolism. The objective of this analysis is to quantify the effect of the initial yeast extract concentration on biomass evolution using the single population model M2.

**Figure 4:**
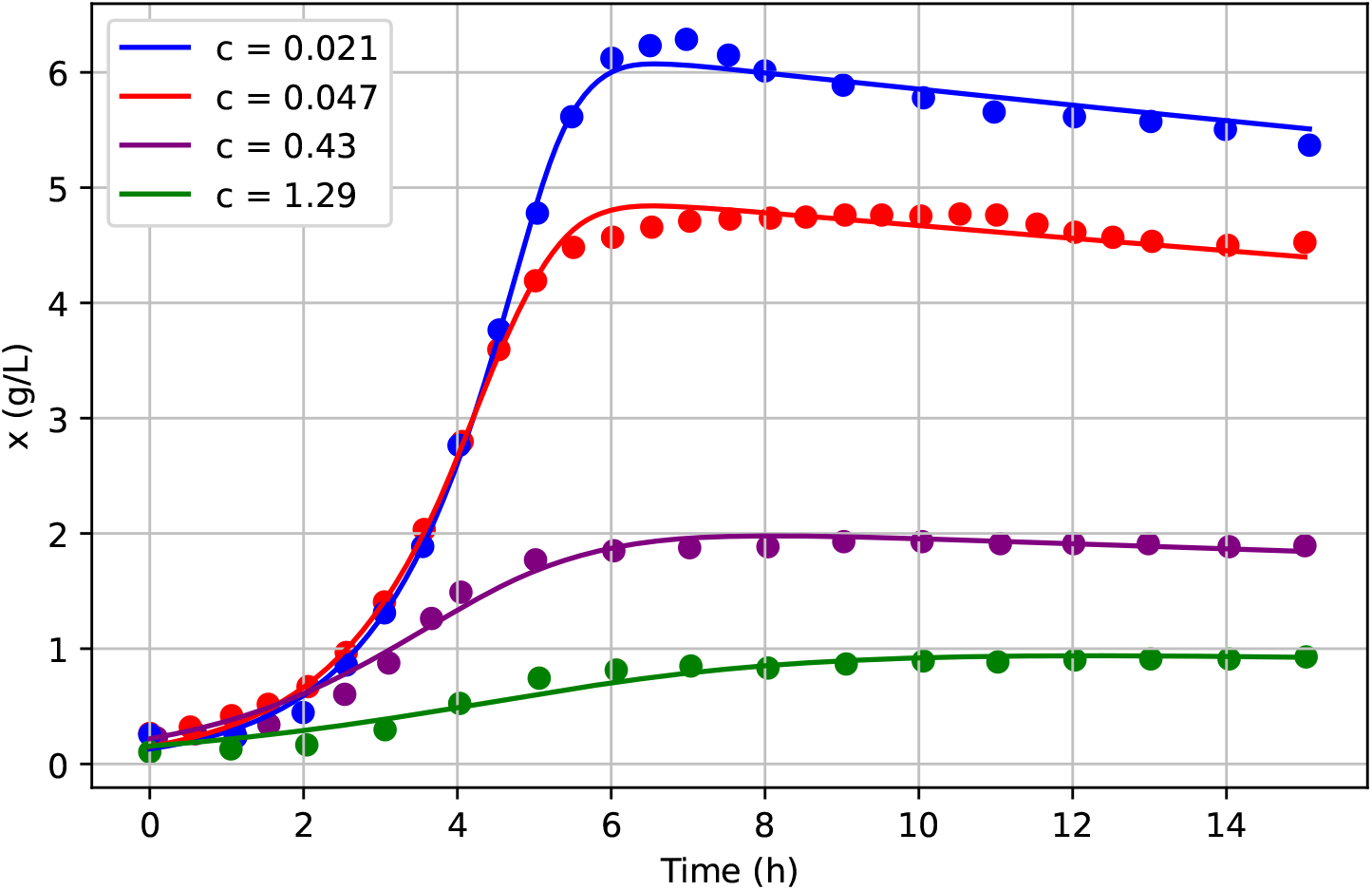
Model M2 fitted to data from [27]. Parameters *a, b*, and *m* are the same across all data sets, while *c* is estimated for each dataset. Parameters are provided in Table 2.

Since the parameter *c* is associated with the initial substrate concentration (see (20)), we hypothesize that the model M2 can describe the data for all conditions by varying only this parameter while keeping *a, b*, and *m* fixed across all the cultivation conditions. This assumption leads to the following vector of parameters to be estimated:

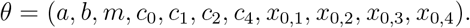

**Table 2:** Parameter values for *c* and *x*_0_ under different initial yeast extrac concentrations. The remaining parameters are fixed at *a* = 0.79 h^−1^, *b* = 0.50 g^−1^ L h^−1^, and *m* = 0.012 h^−1^.

| Yeast extract<br>concentration (g/L) | $c$ | $x_0$ (g/L) |
| --- | --- | --- |
| 20 | 0.021 | 0.130 |
| 10 | 0.047 | 0.152 |
| 5 | 0.430 | 0.219 |
| 2 | 1.290 | 0.155 |

Here, *c*_*j*_ and *x*_0,*j*_ represent the parameter *c* and the initial biomass concentration, respectively, associated with condition *j*. To fit the model to the experimental data, we formulate the following least-squares optimization problem:

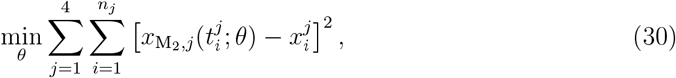

where 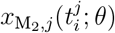 represents the model prediction for condition *j* at time 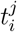, and 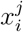 denotes the corresponding experimental measurement. The number of available observations for each condition is given by *n*_*j*_. To solve (30), we employ the Levenberg-Marquardt algorithm using the `curve_fit` function from the `SciPy.optimize` [29] module in Python.

Figure 4 shows the fitting of the model, while Table 2 presents the estimated parameters. The results indicate that conditions with higher initial nutrient concentrations correspond to lower values of *c*, reinforcing its interpretation as a resource availability parameter. This trend aligns with equation (20), where *c* decreases as the initial substrate concentration increases. Furthermore, the model successfully captures the experimental trends across all conditions, as evidenced by the close agreement between the fitted curves and the observed data. These results support the validity of the proposed parameterization and confirm that the model can describe microbial growth dynamics under varying initial conditions with a consistent biological interpretation.

## 5 Conclusions

We have developed and analyzed two novel integro-differential models for single-population dynamics. Unlike classical models such as the logistic growth equation, which only accounts for population stabilization at equilibrium, the proposed models incorporate a decline phase. This feature makes them suitable for studying populations that exhibit a decrease over time, such as those in bioreactor batch cultures, which are widely used in biotechnology and bioprocessing. Through numerical optimization and fitting to experimental data, we demonstrated that these models accurately describe population dynamics observed in mammalian cell and bacterial cultures.

The proposed models provide valuable insights into estimating parameters such as the maintenance rate, a crucial but difficult-to-measure parameter [31], essential for bioprocess control and optimization [32]. This is because our models emerge naturally from fundamental batch reactor equations, preserving biological interpretability. Each parameter retains a clear mechanistic meaning, allowing for more insightful data interpretation and parameter estimation. Indeed, we highlighted that estimating all parameters of a traditional batch culture model solely from population data is often infeasible due to overparameterization and data limitations. However, the structure of our single-population models allows for parameter estimation that is not only mathematically sound but also biologically meaningful.

Beyond batch cultures, the proposed models hold potential for describing any biological system exhibiting a population decline, whether due to resource limitations, environmental constraints, or other factors. Similar to how the logistic growth equation was initially developed for specific scenarios and later found widespread use in diverse fields, the proposed models have the potential for broader applications in ecology and population biology.

Developing single-population models that incorporate the decay phase observed in batch cultures offers several advantages over more complex models that account for multiple state variables. One key advantage is simplified parameter estimation, as these models require fewer parameters, reducing computational costs. In contrast, multi-variable models often suffer from data weighting issues and parameter identifiability problems, complicating their calibration with experimental data [33]. Additionally, their simplicity and broad applicability make them valuable educational tools, providing an intuitive introduction to microbial kinetics before transitioning to more complex modeling frameworks [34].

By addressing key limitations of traditional models and offering a mechanistically grounded alternative, our approach contributes to a deeper understanding of microbial population dynamics in batch bioreactors and beyond.

## Acknowledgments

This study was partly supported by ITMO Cancer of Aviesan within the framework of the 2021-2030 Cancer Control Strategy and by the region Provence Alpes Côte d’Azur with the fellowship “Jeune Docteur Innovant”.

## Data availability statement

The datasets used for model fitting in this study were obtained from previously published sources. Specifically, the mammalian cell culture data was sourced from [16], while the bacterial batch culture data was taken from [27].

The scripts used for model fitting and for generating Figures 3 and 4 are available at https://github.com/mariellePr/Extended-Logistic-Growth-Model.

## Notes

https://github.com/mariellePr/Extended-Logistic-Growth-Model

### Competing Interest Statement

The authors have declared no competing interest.

